# Context-dependent correlations mislead transcriptomic network inference in bulk and single-cell data

**DOI:** 10.64898/2026.06.23.733936

**Authors:** Amir Asiaee, Polina Bombina, Reginald L. McGee, Jake Reed, Zachary B. Abrams, Lynne V. Abruzzo, Kevin R. Coombes

## Abstract

**Background:** Correlation is the dominant input to co-expression module discovery and miRNA-target inference. Both rely on an implicit assumption: a Pearson coefficient pooled across heterogeneous samples, whether tissues, cancer types, or cell types, estimates one biologically meaningful quantity. Simpson’s paradox makes this assumption fragile in principle, since between-group mean shifts can dominate or reverse within-group associations. How often this happens in real transcriptomic data has not been quantified.

**Results:** Across 8,890 TCGA tumors from 31 cancer cohorts and 23,170,038 miRNA–mRNA pairs, 94.8% of pairs showed both positive and negative within-cohort correlations. Restricting to the high-variance domain of one million pairs, 13.3% of pooled correlations with |*r*_global_|≥0.2 reversed against the within-cohort majority at sign tolerance *ε* = 0.05. Heterogeneity was the rule rather than the exception (median *I*^2^ = 0.86, IQR 0.80–0.90), and 99.5% of pairs rejected equal correlation across cohorts at FDR *<* 0.05. Of 692,770 experimentally validated miRTarBase v10 targets measurable in our data, only 0.9% were uniformly negative across cohorts. The pattern recurred across modalities. In GTEx, 21.0% of pooled signs disagreed with the tissue majority, and 23.5% of pairs flipped sign after tissue-mean removal. In 10x PBMC scRNA-seq, 13.1% of gene–gene correlations flipped after cell-type-mean removal; in CITE-seq, 37.9% of protein–RNA pairs flipped under a joint WNN partition of cells. Refining context reduced reversal, though by how much depended on the partition: within BRCA, 5.5% of pairs reversed under molecular PAM50 subtypes versus 0.35% under clinical IHC receptor status, and refining T cells into transcriptome-defined subtypes cut PBMC reversal from 11.8% to 0.13%.

**Conclusions:** A single pooled correlation coefficient can invert direction relative to its within-context constituents at rates that are not negligible. Correlations should be reported with their context: the within-context distribution, a heterogeneity statistic, and a diagnostic that separates between-context mean shifts from within-context association. We provide a small R interface that computes these summaries.

## Background

Correlation between expression measurements is the primary input to a long line of co-expression and network-inference tools: weighted correlation networks [1], mutual-information regulator inference [2], tree-ensemble regulatory networks [3], and message-passing refinement of predicted edges [4]. In transcriptomic studies, miRNA–mRNA correlation in particular is interpreted in regulatory terms, because miRNAs reduce target abundance, so a negative correlation is often taken as evidence consistent with repression [5]. These methods take a pooled correlation matrix as input and treat each pairwise coefficient as a stable summary of the underlying relationship. Downstream analyses inherit that assumption: edges are ranked, modules are called, and regulators are nominated from coefficients estimated across whichever samples happened to enter the matrix. The implicit claim is that a single number per pair is enough.

Correlation, however, is a population quantity, not a property of a single sample. When samples are pooled from heterogeneous contexts, distinct cancer types, normal tissues, cell types, or measurement modalities, the global correlation can disagree with every within-context correlation it summarizes. This conflation has been recognized in statistics for more than a century: it is the kernel of Simpson’s paradox, in which a marginal association reverses or vanishes after stratification by a relevant covariate [6, 7]. In epidemiology and causal inference, Simpson reversals are a standard motivating example for thinking carefully about confounding and effect modification [8, 9]; in machine-learning theory, the conflation of within- and between-group variation is treated as a primary source of spurious association [10]. Genomics inherits the same arithmetic. A pooled coefficient mixes within-context covariation with between-context mean shifts, and the two components can have opposite signs.

In genomics, the closest established response to between-sample heterogeneity is batch correction, by empirical-Bayes adjustment of nuisance differences [11] or surrogate-variable estimation when those differences are unobserved [12]. Both are designed to remove unwanted variation rather than to characterize biologically meaningful heterogeneity, and they leave correlations between features within a sample mostly untouched. A separate line of work on differential co-expression and condition-specific networks asks how pairwise associations change between groups [13], but those methods typically compare two precomputed networks rather than quantify how often a single pooled coefficient misrepresents its constituent groups. At single-cell resolution, co-expression replicability is markedly higher within than across cell types [14], consistent with the broader concern: pooling across contexts can wash out, or invert, structure that is clean inside each context.

None of this prior work answers the empirical question that should set the prior on any pooled correlation in modern genomics: across the kinds of datasets analysts actually use, how often does a pooled correlation disagree with the within-context relationship it is taken to summarize, and how does this depend on how contexts are defined? The technical machinery for answering this question, meta-analytic heterogeneity statistics (Cochran’s *Q*, the *I*^2^ index [15]), variance decomposition, and mean-residualization, has been available for decades, but has not been applied to transcriptomic correlation at scale. The miRNA–mRNA case is a useful entry point, because experimentally validated targets [16] give a built-in check on whether the sign of a pooled coefficient tracks known regulation.

We quantify pooled-versus-within-context disagreement across four independent settings, each pairing a correlation of interest with the context it is pooled across: miRNA–mRNA correlations in TCGA tumors, pooled across cancer types (tissue of origin) [17, 18]; mRNA–mRNA correlations in GTEx, pooled across normal tissues [19]; gene–gene correlations in single-cell RNA-seq of peripheral blood mononuclear cells (PBMC), pooled across cell types [20]; and protein–RNA correlations in CITE-seq, where surface protein and mRNA are measured in the same cells, pooled across cell types defined by weighted nearest-neighbor integration of the two modalities [21, 22]. Across all four, pooled correlations frequently invert the within-context relationship; the effect is not confined to weak associations, persists under standard meta-analytic diagnostics, and depends on how context is defined. Within breast cancer we further compare molecular (PAM50) and clinical (IHC) partitions to show that the choice of context, not just its presence, changes the answer. From these results we suggest a five-item reporting checklist that turns context dependence from a hidden confound into a reported quantity, implemented as a lightweight R interface compatible with standard co-expression workflows. The remainder of the paper develops these results in turn.

## Results

### Pooled miRNA–mRNA correlations conflate within- and between-cohort structure

We computed the global Pearson *r* (*r*_global_) across the 8,890 TCGA samples and within each of the 31 cohorts (*r*_*c*_), following the preprocessing pipeline of [17, 18]. The global distribution is sharply centered near zero with long tails (Fig. 1A). Extreme pooled values do not imply consistent within-cohort relationships: a large global *r*_global_ conflates within-cohort covariance with between-cohort mean shifts. The liver-enriched miRNA miR-122 [23] makes this concrete (Fig. 1B). For the two miR-122 pairs with the most extreme pooled correlations (vs *CFHR2, r*_global_ = 0.97; vs *SLC25A36, r*_global_ = −0.73), most cohorts sit near zero or have miR-122 below detection. The extreme pooled value reflects which cancer types express miR-122, not a within-cohort relationship.

**Figure 1:**
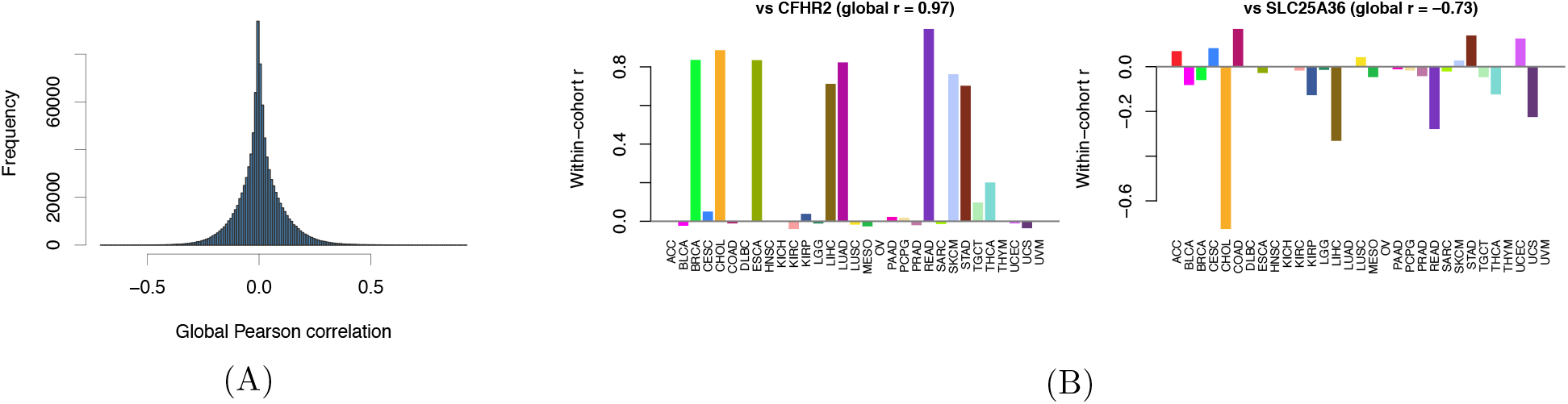
Pooled pan-cancer miRNA–mRNA correlations conflate within- and between- cohort structure. **(A)** Distribution of pan-cancer Pearson *r* across 23,170,038 miRNA–mRNA pairs in TCGA, centered near zero with long tails. **(B)** Within-cohort *r* for miR-122 against two pairs whose pan-cancer values sit at the extremes: *CFHR2* (*r*_global_ = 0.97) and *SLC25A36* (*r*_global_ = −0.73). Within-cohort values cluster near zero or are undefined in cohorts where miR-122 is not expressed.

Genome-wide, 94.8% of the 23,170,038 pairs exhibit both positive and negative correlations across cohorts. For the vast majority of pairs, “the” correlation is not a single stable quantity. Moreover, this sign disagreement does not merely reflect small, noisy per-cohort correlations scattered around zero. To see this, we restrict attention to high-variance pairs (the 10^6^ pairs among the 5,000 most variable mRNAs and 200 most variable miRNAs): even here, 99.5% of pairs that reach |*r*_*c*_| ≥ 0.2 in at least one cohort still split in sign across cohorts, indicating genuine, cohort-specific structure. This also explains why neither standard way of prioritizing a pair from the pooled matrix, by statistical significance or by correlation magnitude, is reliable. A small *r*_global_ can be statistically significant simply because *n* = 8,890 is large, and a large |*r*_global_| can come from between-cohort composition (as with miR-122) rather than a relationship that holds within cohorts.

### A worked Simpson reversal in TCGA: miR-200c-3p versus *HTRA3*

The pair miR-200c-3p vs *HTRA3* is representative (Fig. 2). The pooled scatter is positive (Fig. 2A, *r*_global_ = 0.403), yet the within-cohort *r*_*c*_ is negative in 22 of 31 cancer cohorts (Fig. 2B). Decomposing the pooled covariance separates the two contributions. The between-cohort component is positive: cohort means of miR-200c-3p and *HTRA3* covary positively (Fig. 2C, *r*_means_ = 0.515). The within-cohort component is negative.

**Figure 2:**
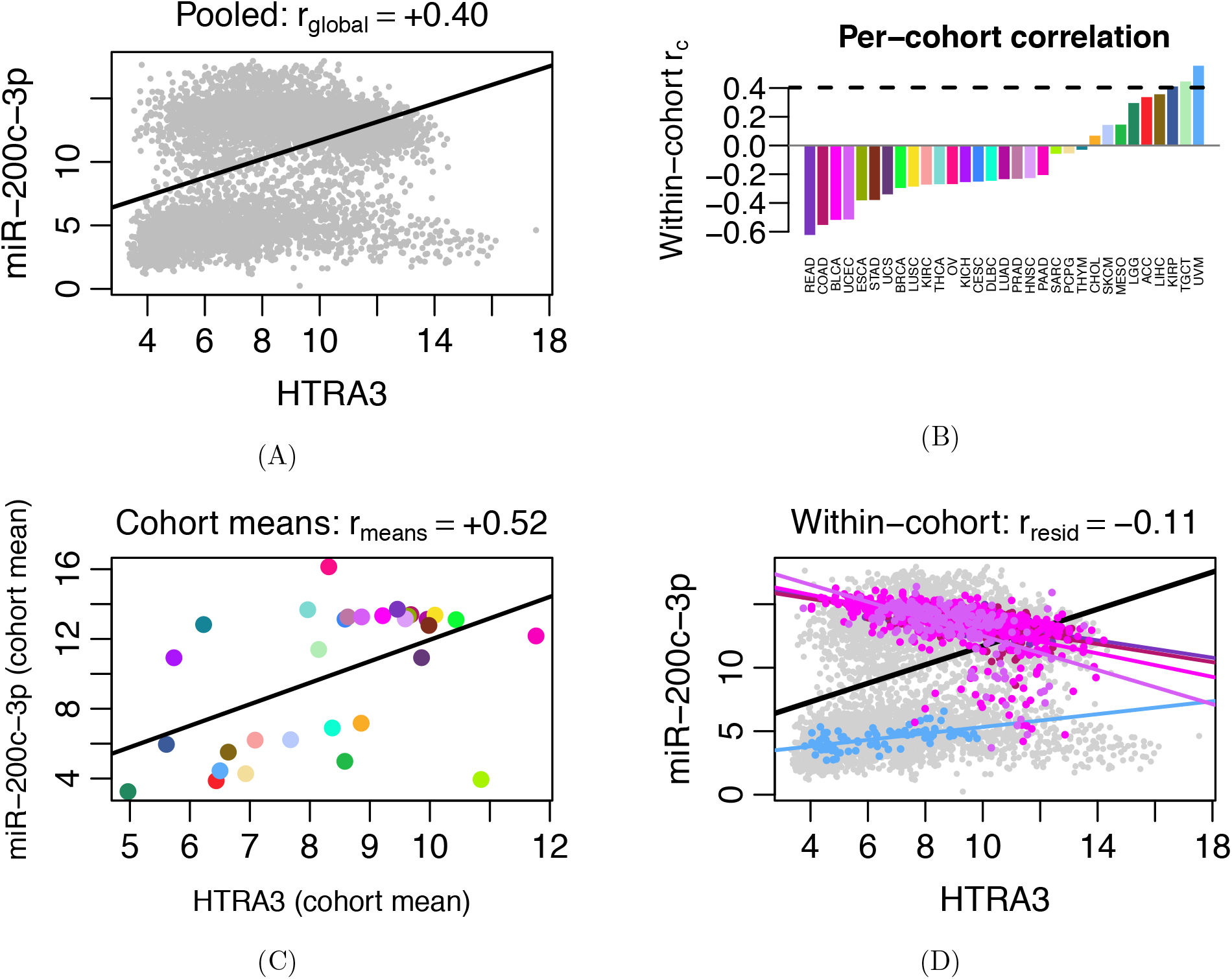
A worked Simpson reversal in TCGA: miR-200c-3p versus *HTRA3*. **(A)** Pan-cancer scatter with a single pooled regression line, *r*_global_ = 0.403. **(B)** Within-cohort *r*_*c*_ across the 31 TCGA cohorts; the association is negative in 22 of 31. **(C)** Cohort-mean scatter of mean(miR-200c-3p) against mean(*HTRA3*), *r*_means_ = 0.515, showing that the pooled positive trend is carried by between-cohort means. **(D)** Pooled scatter with one regression line per cohort overlaid; the within-cohort slopes run opposite to the pooled line, with mean-residualized *r*_resid_ = −0.110 and mixed-effects population slope 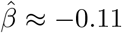.

Per-cohort regression lines on the pooled data run opposite to the global regression line (Fig. 2D), the quantitative signature of Simpson’s paradox [7]. Mean-residualization, which subtracts each cohort’s mean from both features before correlating, turns the pooled positive association negative (*r*_resid_ = −0.110), and a linear mixed-effects model with random cohort slopes recovers a negative population slope 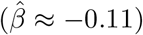 with substantial between-cohort variability. An analyst reading only the pooled *r*_global_ would assign the wrong direction of association. We summarize this pattern across pairs with two counts. To avoid treating near-zero noise as a direction, we score a cohort’s correlation as positive when *r*_*c*_ ≥ *ε* and negative when *r*_*c*_ ≤ −*ε*, using a sign tolerance *ε* = 0.05 unless stated otherwise. A pair is *mixed-sign* when both signs occur among its cohorts, and a *Simpson reversal* when its pooled *r*_global_ is nonzero but takes the opposite sign to the cohort majority; full definitions are in Methods.

### Genome-wide heterogeneity and validated targets

Both the mixed-sign fraction and the Simpson-reversal fraction persist as the effect-size threshold tightens (Fig. 3A,B). In the high-variance domain, 13.3% of pairs with |*r*_global_| ≥ 0.2 reverse against the within-cohort majority at the sign tolerance *ε* = 0.05, and the curves are nearly flat as the threshold or *ε* rise. Context dependence is not an artifact of weak effects.

**Figure 3:**
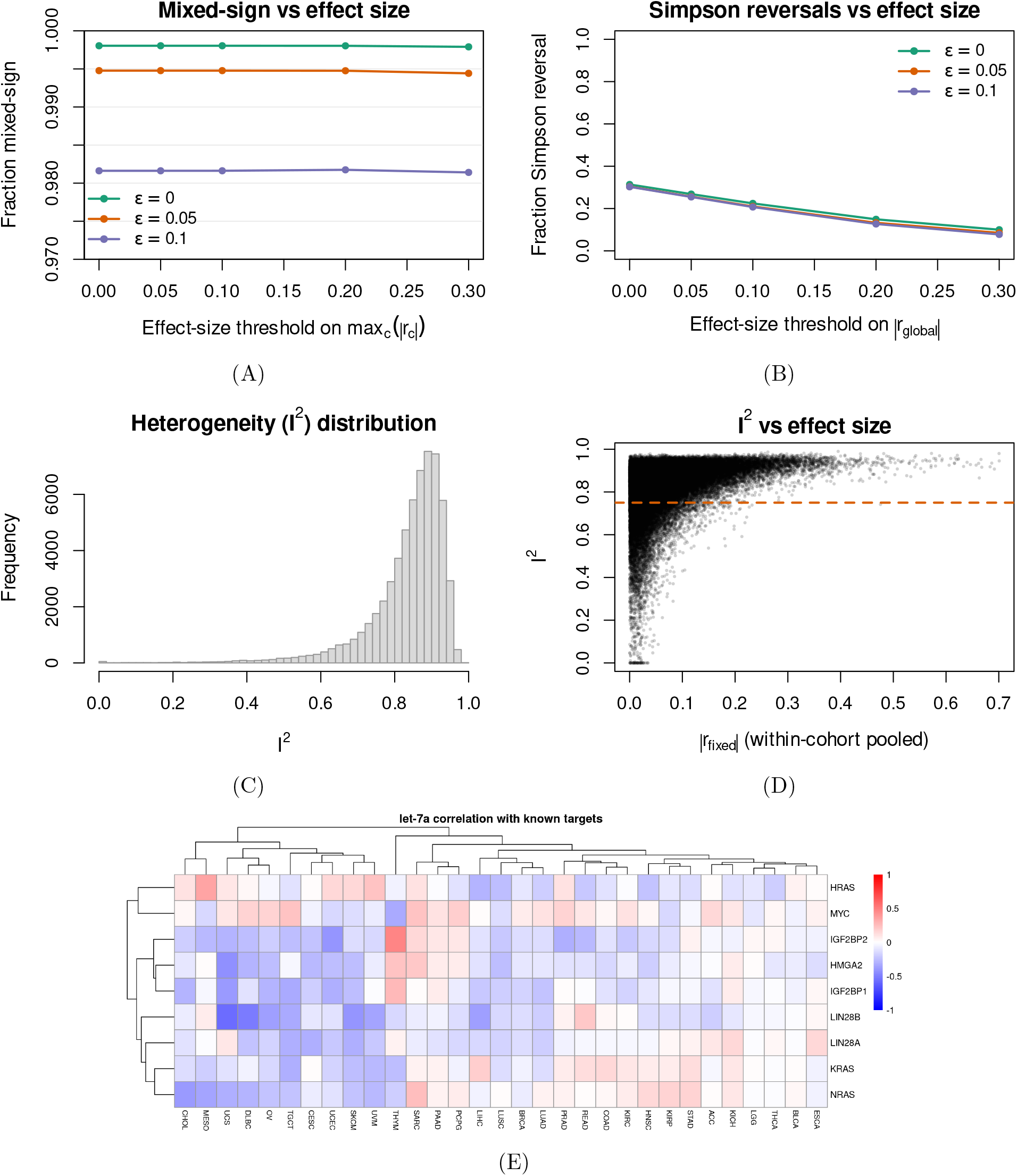
Genome-wide heterogeneity and reversal among validated miRNA targets in TCGA. **(A)** Fraction of mixed-sign pairs as a function of the effect-size threshold max_*c*_ |*r*_*c*_| at sign tolerances *ε* ∈ {0, 0.05, 0.10}. **(B)** Simpson-reversal fraction on the same axes; at *ε* = 0.05, 13.3% of pairs with |*r*_global_| ≥ 0.2 reverse against the within-cohort majority. **(C)** Distribution of *I*^2^ across the 10^6^ high-variance pairs (top 5,000 mRNAs × top 200 miRNAs), median *I*^2^ = 0.86, IQR 0.80–0.90. **(D)** *I*^2^ against the fixed-effects pooled within-cohort correlation |*r*_fixed_| (Methods), showing high heterogeneity across the range. **(E)** Heatmap of within-cohort Pearson *r*_*c*_ for let-7a against its miRTarBase v10 validated targets (rows) across TCGA cohorts (columns); even canonical let-7a targets span −0.6 to +0.5.

Formal meta-analysis agrees with these observations. Correlation heterogeneity across cohorts is extreme (median *I*^2^ = 0.86, IQR 0.80–0.90; Fig. 3C) and persists across effect sizes (Fig. 3D). Cochran’s *Q* rejects equal correlation across cohorts for 99.5% of pairs at FDR *<* 0.05 [24]. For the vast majority of pairs in the high-variance domain, no single within-cohort *r* is compatible with all 31 cohorts simultaneously.

If negative miRNA–mRNA correlation were a reliable proxy for repression, miRTarBase-validated targets [16] should be enriched for consistently negative within-cohort correlations. They are not (Fig. 3E). Across the 692,770 validated target pairs measurable in our data, only 6,413 (0.9%) are uniformly negative across cohorts. Canonical let-7a targets including the RAS family, *HMGA2*, and *LIN28A*/*B* span −0.6 to +0.5 across cancer types, and several are frequently positive. Experimental validation therefore does not guarantee a consistent correlation: it confirms that the interaction can occur, not that its sign is stable across cancer types. A target relationship that holds in one tumor type can invert in another, so reporting a single pooled *r* for a validated pair will, on average, give a direction-of-association estimate that contradicts most of the underlying tissues.

### Context dependence in GTEx mRNA–mRNA correlations

The phenomenon is not cancer-specific (Fig. 4). In GTEx mRNA–mRNA correlations across the 20 largest tissues (200,000 pairs sampled from the 1,500 most variable genes; tissue assigned from the SMTSD field of the GTEx v8 annotations) [19], the median *I*^2^ across pairs is 0.71. 21.0% of pooled correlations disagree in sign with the tissue majority (6.2% among pairs with |*r*_global_| ≥ 0.2), and 23.5% flip sign after subtracting tissue means.

**Figure 4:**
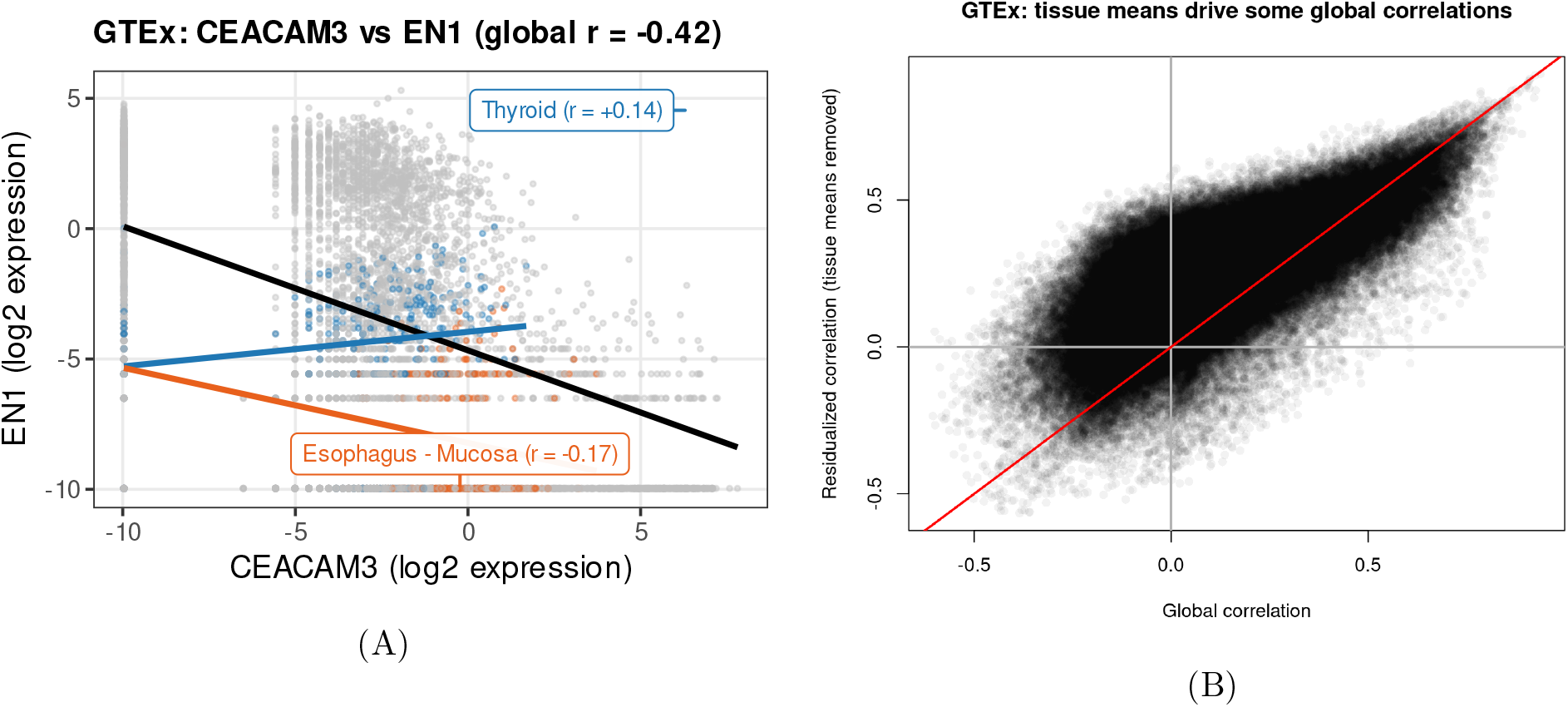
Tissue context reverses mRNA–mRNA correlations in GTEx. **(A)** *CEACAM3* against *EN1* across 20 GTEx tissues (≥ 150 samples each); the pooled regression line gives *r*_global_ = −0.42, while within-tissue slopes are positive in 15 of 20 tissues. **(B)** Pooled *r*_global_ against tissue-mean-residualized *r*_resid_ for 200,000 sampled pairs from the 1,500 most variable genes; 23.5% of pairs flip sign after tissue-mean removal, and 6.2% reverse among pairs with |*r*_global_| ≥ 0.2 (median *I* = 0.71).

The pair *CEACAM3* vs *EN1* illustrates the pattern (Fig. 4A). A negative pooled regression line (*r*_global_ = −0.42) runs opposite to within-tissue regression lines that are positive in 15 of 20 tissues. Aggregated across pairs (Fig. 4B), pooled and tissue-mean-residualized correlations frequently disagree, and a non-trivial cloud of pairs that look strong in the pooled correlation are weak, or oppositely signed, within tissues. Cancer-specific between-cohort structure is therefore one instance of a broader pattern that also operates across normal tissues.

### Reversal in single-cell RNA-seq and CITE-seq

Bulk results could in principle be driven by tissue composition, so we repeated the analysis in single-cell data. Marker-based cell-type labels are assigned before the correlation analysis, so the context definition is independent of the pair-specific correlation being summarized. In PBMC scRNA-seq, 13.1% of global gene–gene correlations flip sign after cell-type-mean removal. The pair *LYZ* vs *FTH1* illustrates this (Fig. 5A): a positive pooled correlation (*r*_global_ = 0.68) is driven almost entirely by between-cell-type composition (cell-type-means *r* = 0.99) and collapses to *r*_resid_ = −0.04 within cell type. Single-cell co-expression has separately been observed to be substantially more replicable within than across cell types [14].

**Figure 5:**
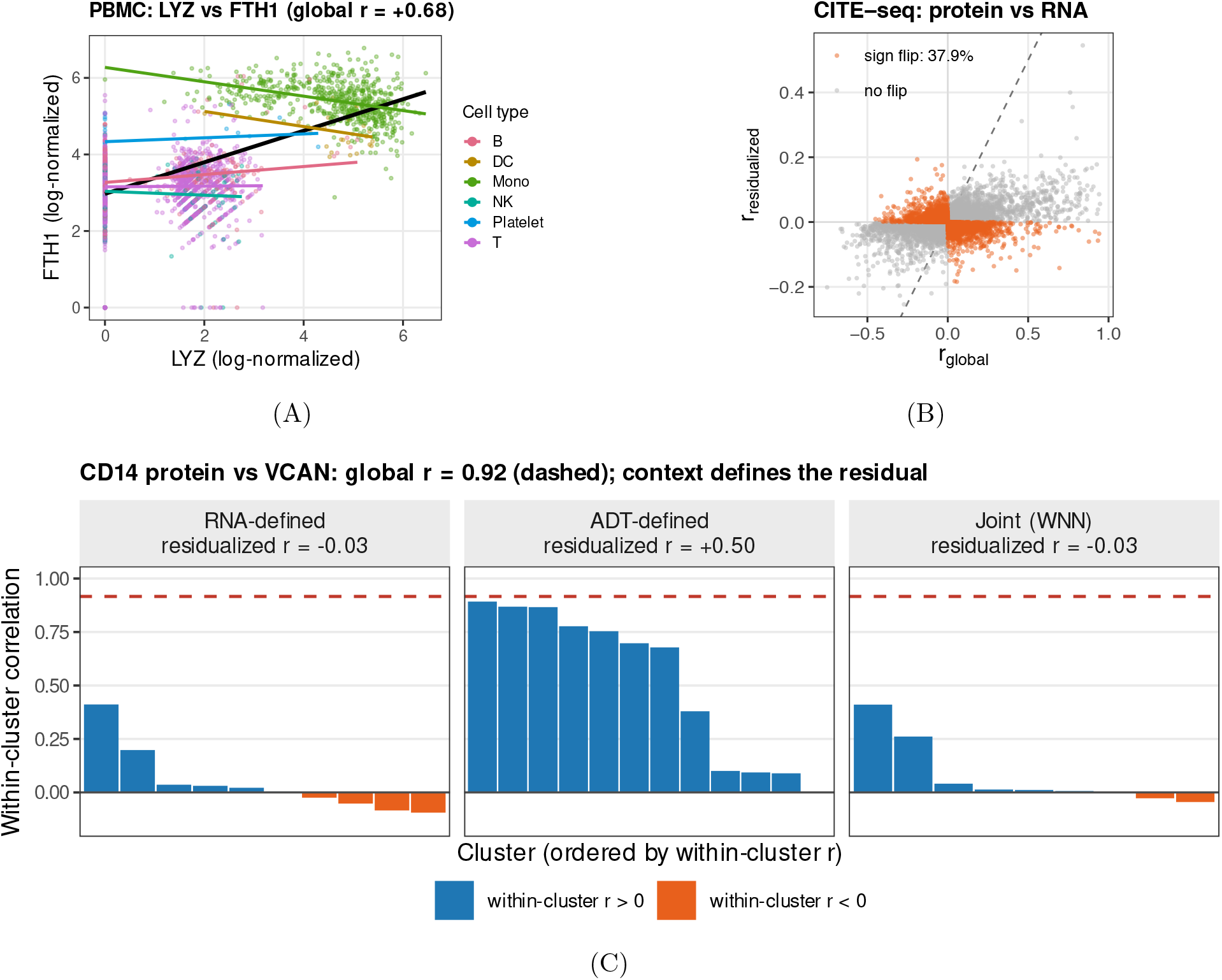
Reversal in single-cell RNA-seq and CITE-seq depends on the chosen context. **(A)** PBMC scRNA-seq (10x Genomics PBMC 3k, ∼2,700 cells): *LYZ* against *FTH1* across six coarse cell types (T, B, NK, monocytes, dendritic, platelets), *r*_global_ = 0.68 driven by cell-type means (*r* = 0.99); cell-type-mean-residualized *r* = −0.04. **(B)** CITE-seq protein–RNA pairs (7,798 cells, 14,770 pairs), pooled *r*_global_ against joint-WNN-residualized *r*_resid_; 37.9% of pairs flip sign.**(C)** CITE-seq exemplar CD14 surface protein, measured by antibody-derived tag (ADT), against *VCAN* mRNA under three partitions: *r*_resid_ = −0.03 for RNA-defined Louvain clusters, 0.50 for ADT-defined clustering, and −0.03 for joint WNN, while *r*_global_ = 0.92.

The same compositional mechanism extends to multi-modal CITE-seq [21], where surface protein measured by antibody-derived tag (ADT) and mRNA are measured in the same 7,798 cells. A cell’s context here can be defined in three ways: by clustering on RNA alone (Louvain clustering of the transcriptome), on ADT alone, or on a joint representation that integrates both modalities through weighted nearest-neighbor (WNN) analysis [22]. Under the joint WNN partition, 37.9% of protein–RNA pairs flip sign after mean-residualization (Fig. 5B). The definition matters: for CD14 protein vs *VCAN* mRNA (Fig. 5C), the pooled correlation is large and identical under all three partitions (*r*_global_ = 0.92, because pooling ignores cluster labels), yet the residualized correlation swings with the partition, from *r*_resid_ = −0.03 under RNA-based Louvain clustering to 0.50 under ADT-based clustering and back to −0.03 under WNN. The chosen partition decides whether the association is read as compositional or as genuinely within-context.

### Refining context within breast cancer attenuates reversal

Reversal generally recedes as broad contexts are refined toward biologically coherent strata (Fig. 6), but how much it recedes depends on how those strata are defined. Restricting matched TCGA samples to BRCA tumors and joining sample-level subtype labels from the UCSC Xena BRCA clinical matrix [25], we recompute within-subtype correlations among mRNAs and miRNAs on a BRCA-specific high-variance domain under two alternative subtype definitions: a molecular one, PAM50 (LumA, LumB, Basal, Her2, Normal-like; *n* = 510), and a clinical one, IHC receptor status (HR+, HER2+, TNBC; *n* = 403).

**Figure 6:**
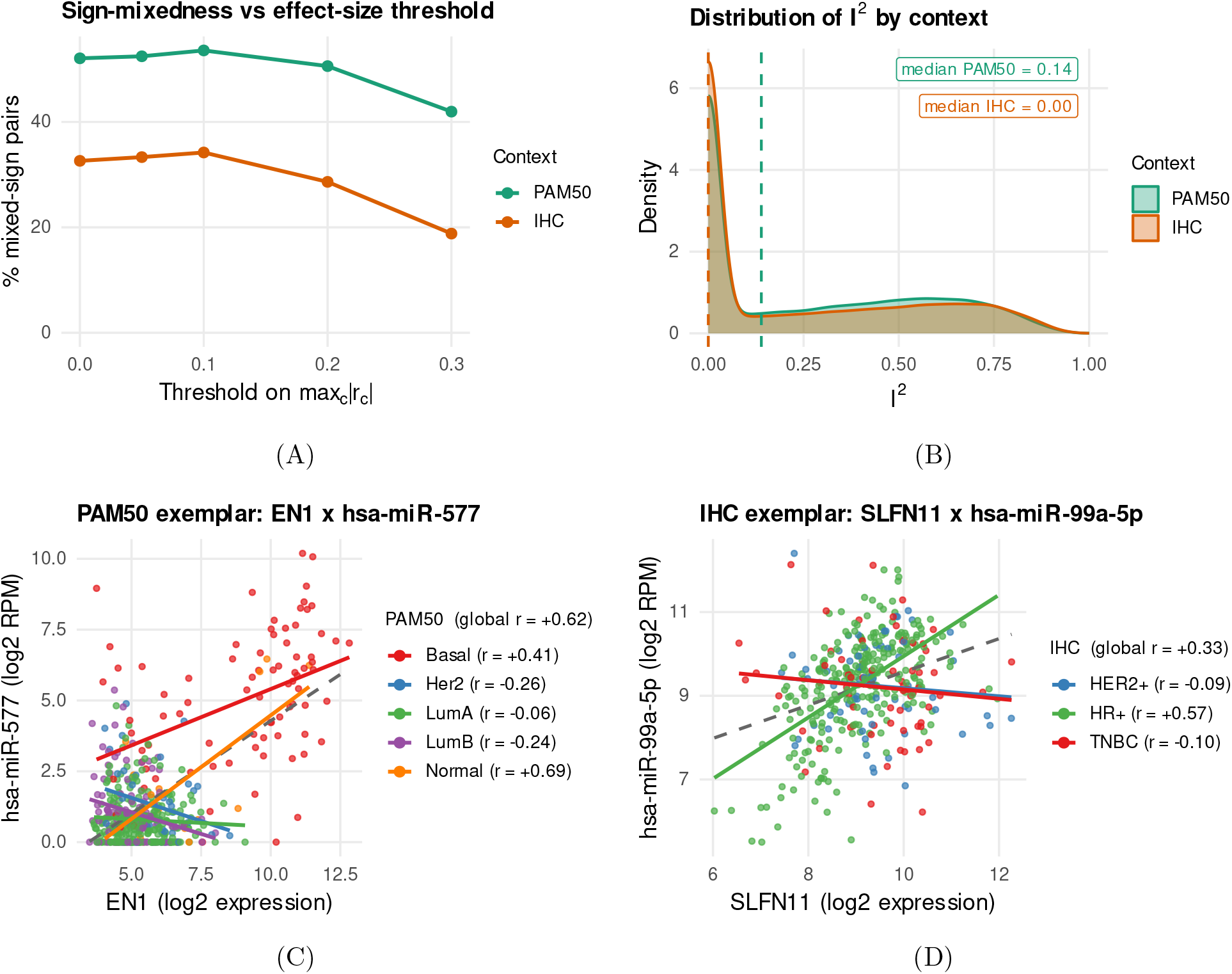
Refining context within breast cancer collapses heterogeneity under IHC but not PAM50. Partitions compared are the five PAM50 intrinsic subtypes (LumA, LumB, Basal, Her2, Normal-like; *n* = 510 tumors) and immunohistochemistry (IHC) receptor groups HR+ (hormone-receptor-positive), HER2+ (HER2-amplified), and TNBC (triple-negative breast cancer; *n* = 403). **(A)** Within-BRCA mixed-sign fraction as a function of effect-size threshold under PAM50 versus IHC; at max_*c*_ |*r*_*c*_| ≥ 0.2, 50.6% remain mixed-sign under PAM50 against 28.6% under IHC. **(B)** *I* distribution by partition, median 0.14 under PAM50 versus 0.00 under IHC. **(C)** PAM50 exemplar *EN1* against hsa-miR-577, *r*_global_ = +0.62 with per-subtype *r* ranging from −0.26 to +0.41. **(D)** IHC exemplar *SLFN11* against hsa-miR-99a-5p, *r* = +0.57 in HR+ tumors and near zero or negative in HER2+ and TNBC.

Under PAM50 (Fig. 6A,B), 50.6% of pairs with max_*c*_ |*r*_*c*_| ≥ 0.2 remain mixed-sign and 5.5% of pairs with |*r*_global_| ≥ 0.2 show Simpson reversal, with median *I*^2^ ≈ 0.14. Under IHC (Fig. 6A,B), the figures are 28.6% mixed-sign and 0.35% reversal, with median *I*^2^ ≈ 0. Both fall well below the cross-cancer rate, so refining a pan-cancer pool into BRCA subtypes attenuates reversal under either definition; but the molecular and clinical cuts disagree about how much heterogeneity remains.

The exemplars make the contrast concrete. *EN1* vs hsa-miR-577 has *r*_global_ = +0.62 yet per-subtype correlations range from −0.26 to +0.41 under PAM50 (Fig. 6C). *SLFN11* vs hsa-miR-99a-5p has *r* = +0.57 in HR+ tumors but is near zero or negative in HER2+ and TNBC under IHC (Fig. 6D). The same logic applies in single cells: refining T cells into transcriptome-defined subtypes instead of cell-type markers drops the PBMC reversal rate from 11.8% to 0.13% at |*r*_global_| ≥ 0.05, and from 10.3% to 0% at |*r*_global_| ≥ 0.2 (zero of 41 eligible pairs). Refinement therefore helps, but how one refines matters: the chosen partition, not merely its granularity, determines which heterogeneity it resolves and which remains visible. A correlation that survives refinement is more credible as a context-stable descriptive association.

### A reporting checklist for correlation in context

For any prioritized correlation, we recommend reporting five quantities together rather than the pooled coefficient alone:

1. the pooled global correlation across all samples;
2. the full distribution of within-context correlations, with context sample sizes;
3. heterogeneity statistics, Cochran’s *Q* with FDR control and the *I*^2^ index [15];
4. a pooled within-context estimate, random-effects when heterogeneity is substantial (e.g. *I*^2^ *>* 0.75) [26];
5. at least one diagnostic that distinguishes between-context mean shifts from within-context association, such as mean-residualization or a mixed-effects model.

These quantities are lightweight to compute from any context-annotated expression matrix, can be carried alongside a standard co-expression analysis, and produce interpretable outputs. We provide a small R interface implementing them at https://github.com/AsiaeeLab/context-corr.

## Discussion

### Context heterogeneity is the default, not the exception

Across bulk tumor, bulk normal tissue, single-cell RNA, and paired protein–RNA measurements, pooled pairwise correlations frequently disagree with their within-context constituents, and the disagreement is structural. It survives effect-size filtering, recurs at every threshold we examined, is captured by standard meta-analytic heterogeneity, and arises through a compositional mechanism: the pooled coefficient absorbs between-context shifts in the two features’ mean levels, and the distortion grows as those mean shifts grow. The pattern does not localize to a particular cohort, modality, or correlation cutoff.

A strong pooled correlation carries an implicit homogeneity assumption that, for transcriptomic features measured across distinct cancer types, tissues, or cell types, is rarely satisfied in practice. When samples are drawn from a mixture of contexts with appreciably different means, the pooled coefficient summarizes a quantity that may not exist within any single context. The right baseline expectation when pooling is heterogeneity. Consistency across contexts is what should require evidence.

### Relation to co-expression and causal-inference literature

Methods that take a pooled correlation matrix as input (WGCNA [1], ARACNe [2], GENIE3 [3], and PANDA [4]) implicitly treat pooled co-expression as the relevant association. Differential and condition-specific co-expression analyses [13] explicitly compare networks across groups, and at single-cell resolution co-expression replicates more cleanly within than across cell types [14]. Our diagnostics sit upstream of all of these. They quantify how often the pooled input is itself misleading, and they apply no matter how the downstream method turns those associations into edges: by hard-thresholding correlations into an unweighted network, raising them to a soft power (as in WGCNA), or replacing them with tree-ensemble importance scores (as in GENIE3).

The framing connects to a long causal-inference tradition. Simpson’s paradox is the canonical illustration that stratification can invert a marginal association [6, 7], and the modern treatment of effect modification in directed acyclic graphs makes precise the conditions under which a within-stratum and a marginal estimand differ in sign [8, 9]. Reversal in our setting is a measurement-scale instance of the same phenomenon, with cancer type or cell type playing the role of the modifier. Standard tools for handling unwanted variation in expression data (empirical-Bayes batch correction and surrogate-variable analysis [12]) adjust for nuisance heterogeneity but were not designed to characterize biologically meaningful context heterogeneity. Reversal is precisely what they leave on the table.

## Limitations

The analysis is descriptive. A within-context correlation is not, by itself, a causal claim about within-context regulation; it is only a less confounded summary than its pooled counterpart, and turning a context-stable correlation into a causal claim still requires perturbation or mechanistic follow-up.

We focus on Pearson correlation. A Pearson-versus-Spearman comparison on a high-variance subset (Additional file 1: Fig. S1) shows that the reversal rate is essentially unchanged with Spearman. Bulk TCGA measurements aggregate over cell types within a tumor, so within-cohort correlation still mixes a residual compositional component that our diagnostics cannot fully resolve. The PBMC analysis uses a single donor, so between-donor heterogeneity is outside its scope. CITE-seq partition choice interacts with ADT quality, and our reversal counts depend on whether sparse or noisy ADTs are included.

Context definition is itself a modeling choice. Over-refining contexts when per-context sample sizes are small inflates noise; the framework makes this visible because the reporting checklist lists the per-context sample sizes and the heterogeneity statistics weight each context by its precision, so an under-powered partition is flagged rather than mistaken for genuinely reduced reversal.

### Practical recommendations

The reporting checklist we propose is operational, not philosophical. For any prioritized pair (a candidate co-expression edge, a candidate miRNA-target pair, or a candidate marker-gene relationship), the pooled coefficient, the within-context distribution of *r*_*c*_, the heterogeneity statistic *I*^2^, a random-effects pooled estimate [26], and at least one between-versus-within diagnostic together cost only a few additional lines of code beyond cor(). They keep the pooled summary while making its interpretation explicit. A correlation that survives such refinement is more credible as a context-stable descriptive association; one that does not survives only as a between-context artifact, and should be reported as such. The small R interface that implements these summaries is intended to be used directly by analysts running standard co-expression workflows on context-annotated matrices.

## Conclusions

Pooling miRNA–mRNA correlations across heterogeneous samples can produce coefficients that disagree in sign or magnitude with most of the within-context values that generated them. The pattern holds in bulk TCGA tumors partitioned by cohort, in GTEx tissues, and in single cells partitioned by cell type or by ADT-defined protein bins, and it survives sign-tolerance controls, the choice of correlation coefficient, and changes to the partition. The divergence is structural: it tracks the chosen grouping rather than residual noise, and a prioritized pair from one partition can lose its rank, change sign, or disappear under another. For pair-level inference in mixed-context data, a pooled coefficient is not enough. Reporting the within-context distribution, a heterogeneity summary such as *I*^2^, and a between-versus-within diagnostic alongside the pooled value gives readers the information needed to judge a candidate pair. The accompanying R interface computes these quantities from a single call.

## Methods

### Data sources

The results presented here are based in whole or part upon data generated by The Cancer Genome Atlas (TCGA) Research Network (https://www.cancer.gov/tcga). We used the FireBrowse portal (http://firebrowse.org/) to identify and download matched mRNA-seq and miRNA-seq expression data following the FireBrowse standardized-data release of 28 January 2016. mRNA expression was downloaded per cohort as upper-quartile-normalized RSEM gene-level counts, and miRNA expression was downloaded per cohort as reads-per-million (RPM). Patients were included when both data types were available. We excluded GBM and LAML due to limited matched miRNA-seq in these cohorts; the resulting matched dataset contained 8,890 samples across 31 cancer cohorts: ACC, BLCA, BRCA, CESC, CHOL, COAD, DLBC, ESCA, HNSC, KICH, KIRC, KIRP, LGG, LIHC, LUAD, LUSC, MESO, OV, PAAD, PCPG, PRAD, READ, SARC, SKCM, STAD, TGCT, THCA, THYM, UCEC, UCS, and UVM. Our TCGA download and preprocessing scripts follow procedures described in prior work [17, 18].

For non-cancer validation, we analyzed GTEx mRNA expression across tissues [19]. We used a UCSC Xena Browser expression matrix (“TCGA TARGET GTEx” cohort) [25, 27] with values in log_2_(TPM + 0.001). Tissue of origin was obtained from the detailed tissue-site (SMTSD) field of the GTEx v8 sample annotations, joined to the expression matrix on sample identifier, and used as the context for within-group correlations; the 20 largest tissues (≥ 150 samples each) were retained. Validated miRNA–target relationships were downloaded from miRTarBase version 10.0 [16].

### Single-cell data and preprocessing

#### PBMC scRNA-seq

We used the 10x Genomics PBMC 3k dataset (approximately 2,700 PBMCs from a healthy donor). Preprocessing was performed with Seurat v5 [28]: minimum 200 detected features per cell, maximum 20% mitochondrial reads, log-normalization, top 2,000 highly variable genes for downstream analysis. Coarse cell-type labels (T cells, B cells, NK cells, monocytes, dendritic cells, and platelets) were assigned from established surface-marker genes; these labels were assigned before the pairwise correlation analysis and do not use the tested pair’s correlation value. T cells were re-clustered to identify subtypes for the finer-grained analysis.

#### CITE-seq

We used a PBMC CITE-seq dataset [21] with paired RNA and surface-protein measurements (7,798 cells passing the same quality-control filters). Antibody-derived tag (ADT) counts were normalized using centered log-ratio (CLR) transformation. We defined three alternative contexts by clustering cells using: (1) RNA expression alone (Louvain clustering on the top 20 RNA principal components), (2) ADT expression alone (clustering on CLR-normalized protein levels), and (3) a joint representation via weighted nearest neighbor (WNN) analysis [22]. Correlations were computed between ADT features and RNA features under each context (14,770 protein–RNA pairs per context definition).

### Correlation analysis

We computed correlations on log-transformed expression values. For TCGA mRNA-seq, log_2_(10 + *x*) on FireBrowse RSEM normalized counts. For TCGA miRNA-seq RPM, log_2_(1 + *x*). For GTEx, values were already log_2_(TPM + 0.001). Global correlations were computed across all samples; within-context correlations were computed within each cancer cohort (TCGA), tissue (GTEx), or cell cluster (single-cell). A pair was treated as “measurable” in a context if both variables had non-zero variance after transformation. Unless otherwise stated, we report Pearson correlation; a Pearson-versus-Spearman sensitivity analysis on a high-variance subset confirms qualitative robustness (Additional file 1: Supplementary notes).

For computationally intensive summaries we restrict to a high-variance domain: the top 5,000 mRNAs and top 200 miRNAs by global variance, yielding 10^6^ miRNA–mRNA pairs. For GTEx, the analogous domain comprised the 1,500 most variable genes, from which 200,000 gene–gene pairs were sampled.

### Statistical testing and heterogeneity

The Fisher Z-transformation is 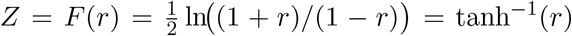; under bivariate normality, *Z* ∼ N (*F* (*ρ*), 1*/*(*N* − 3)) [29].

Heterogeneity across contexts was tested with Cochran’s *Q* statistic

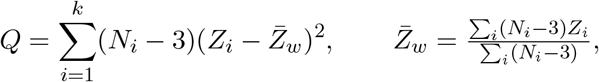

which under the null follows 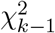. We report the *I*^2^ index [15]:

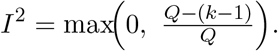

For large-scale summaries we use Benjamini–Hochberg FDR [24]. We compute fixed-effects pooled correlations as 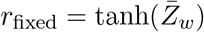 and use |*r*_fixed_| as a within-cohort effect-size summary. For random-effects pooling we use the DerSimonian–Laird estimator [26]. Writing the inverse-variance weights as *w*_*i*_ = *N*_*i*_ − 3 (so 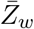 above is the fixed-effects estimate), the between-context variance is estimated by

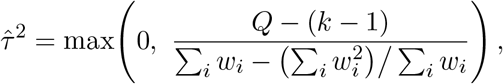

and the random-effects pooled Fisher-*z* reweights each context by 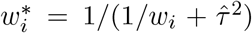, then back-transforms to a correlation:

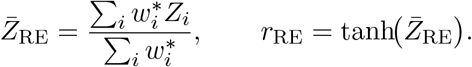

When 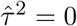 this reduces to the fixed-effects estimate *r*_fixed_; when heterogeneity is present the weights 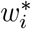are more nearly equal across contexts, pulling the pooled estimate toward the unweighted within-context mean and widening its interval. Random-effects pooled estimates are the within-context summary recommended in the reporting checklist for prioritized pairs.

### Simpson’s paradox diagnostics

For illustrative pairs of molecular features (here a miRNA *M* and an mRNA *G*; the same decomposition applies to any feature pair), we decompose the global covariance into within-context and between-context components,

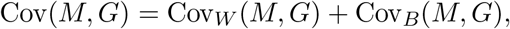

with

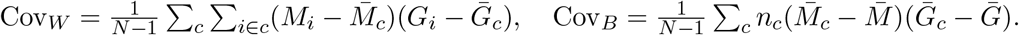

We also compute mean-residualized correlations, cor(*M* ^′^, *G*^′^) with *M* ^′^ = *M* − E[*M* | *C*] and *G*^′^ = *G* − E[*G* | *C*], and fit linear mixed-effects models with random intercepts and slopes by context. In lme4 notation *M* ∼ *G* + (1 + *G* | context), this fits, for sample *i* in context *c*,

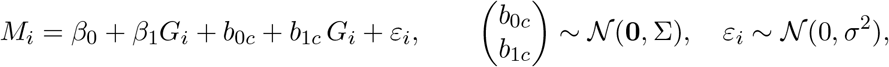

where the fixed slope *β*_1_ is the population-average within-context slope (reported as 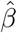 for the worked pair) and the random-effect covariance Σ captures context-to-context variation; its slope variance (Σ_22_) measures how much the within-context slope shifts across contexts.

### Mixed-sign classification and Simpson reversal

Given a sign tolerance *ε* ≥ 0, we define 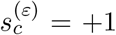 if *r*_*c*_ ≤ −*ε*, and 0 otherwise. A pair is *mixed-sign* if at least one context has *s*_*c*_ = +1 and at least one has *s*_*c*_ = −1. A *Simpson reversal* occurs when 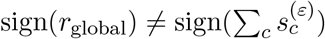 (excluding ties). This majority vote is taken over contexts, not samples; ties and pairs with zero global correlation are excluded. Unless otherwise stated, presented mixed-sign and reversal rates use *ε* = 0.05, and we report their dependence on *ε* ∈ {0, 0.05, 0.1} (Fig. 3A,B).

### BRCA subtype analysis

To test whether Simpson reversal persists at finer biological granularity, we restricted matched TCGA samples to BRCA tumors and joined sample-level intrinsic-subtype labels from the UCSC Xena BRCA clinical matrix [25, 27], matched on the first 15 characters of the TCGA aliquot barcode. Two context definitions were considered: *PAM50* (LumA, LumB, Basal, Her2, Normal-like) from the RNA-seq–based PAM50 subtype call, and *IHC* status combining the estrogen-receptor (ER), progesterone-receptor (PR), and HER2 status annotations into HR+ (ER+ or PR+, HER2−), HER2+ (HER2+ regardless of ER/PR), and triple-negative (ER−, PR−, HER2−). Tumors with equivocal or missing receptor status were excluded, yielding 510 tumors with a PAM50 call and 403 with complete receptor status. The high-variance domain was recomputed within BRCA (top 5,000 mRNAs and top 200 miRNAs by variance among BRCA tumors). All other correlation, heterogeneity, and reversal-rate calculations followed the specifications above.

### Software and code availability

All analyses were performed in R using base R for correlation, with `ggplot2` and `pheatmap` for visualization. Reproducibility scripts and a minimal R interface for “correlation in context” summaries are available at https://github.com/AsiaeeLab/context-corr.

## Declarations

### Ethics approval and consent to participate

Not applicable. This study uses publicly available, de-identified data from TCGA, GTEx, and 10x Genomics public PBMC datasets.

### Consent for publication

Not applicable.

## Availability of data and materials

All datasets analyzed in this study are publicly available. TCGA matched miRNA-seq and mRNA-seq expression data were obtained from FireBrowse (http://firebrowse.org/, “stddata 2016 01 28”). GTEx expression matrices were obtained from UCSC Xena (https://xenabrowser.net/). The PBMC 3k scRNA-seq dataset is publicly distributed by 10x Genomics. The CITE-seq dataset is from Stoeckius et al. [21]. Validated miRNA–target relationships are from miRTarBase v10.0 [16]. All analysis scripts, the minimal “correlation in context” R interface, and processed outputs supporting the conclusions of this article are available at https://github.com/AsiaeeLab/context-corr.

## Authors’ contributions

KRC conceptualized the original study, developed the miRNA–mRNA analysis code, performed the initial analyses, and wrote the original manuscript draft. AA expanded the study, developed and performed the context-dependence and Simpson reversal analyses, extended the analyses across datasets and modalities, generated figures, and revised the manuscript. PB, RLM, JR, ZBA, and LVA contributed to interpretation, provided feedback on experiments, and reviewed and edited the manuscript. All authors read and approved the final manuscript.

## Acknowledgements

The results published here are in whole or part based upon data generated by The Cancer Genome Atlas (TCGA) Research Network. We thank the TCGA, GTEx, and 10x Genomics consortia for making their data publicly available.

## Additional files

### Additional file 1 (PDF, supplement.pdf)

Supplementary material: a glossary of key terms (Table S1), Pearson-versus-Spearman robustness (Fig. S1), single-cell global versus residualized correlations (Fig. S2), T-cell subtype refinement (Fig. S3), and supplementary notes on Pearson-vs- Spearman robustness, rare stable associations, and extended methods.

