## Supplementary Material for "Context-dependent correlations mislead transcriptomic network inference in bulk and single-cell data"

### Additional file 1: Supplementary material

This file provides definitions of key terms, robustness and extension analyses, and extended notes for the Research article.

#### Definitions

Table S1 collects the working definitions used throughout the main text.

Table S1: Key terms used throughout the Research article and this supplement.

| Term | Definition |
|---|---|
| Context | A partition of samples by a biological covariate (e.g., TCGA cancer cohort, GTEx tissue, single-cell cluster, BRCA molecular subtype) within which correlations are computed. |
| Global (pooled) correlation | Pearson correlation $r_{\text{global}}$ computed across all samples, ignoring the context partition. |
| Within-context correlation | Pearson correlation $r_c$ computed using only samples assigned to context $c$. |
| Strong pooled pair | A pair with $\|r_{\text{global}}\| \geq 0.2$, used as the default effect-size threshold for reporting reversal rates. |
| Mixed-sign pair | A pair for which the set $\{r_c\}$ contains both signs after applying the sign tolerance $\varepsilon$. |
| Simpson reversal | A pair for which $\text{sign}(r_{\text{global}})$ disagrees with the within-context majority sign at tolerance $\varepsilon$. |
| Simpson-reversal rate | Fraction of strong pooled pairs that show a Simpson reversal at a given $\varepsilon$. |
| Mean-residualization | Subtraction of context-specific means from each variable before correlation; the resulting Pearson $r_{\text{resid}}$ isolates within-context covariation. |
| Heterogeneity ($Q$, $I^2$) | Cochran's $Q$ and Higgins–Thompson $I^2$ [1] computed on Fisher-Z–transformed $r_c$ values to quantify dispersion across contexts. |
| High-variance domain | Subset of variables retained by variance filtering before pair enumeration (e.g., top 5,000 mRNAs $\times$ top 200 miRNAs in TCGA). |
| Sign tolerance ($\varepsilon$) | A threshold $\|r_c\| < \varepsilon$ that treats a within-context correlation as effectively zero when computing majority sign and mixed-sign status. |

#### Supplementary figures

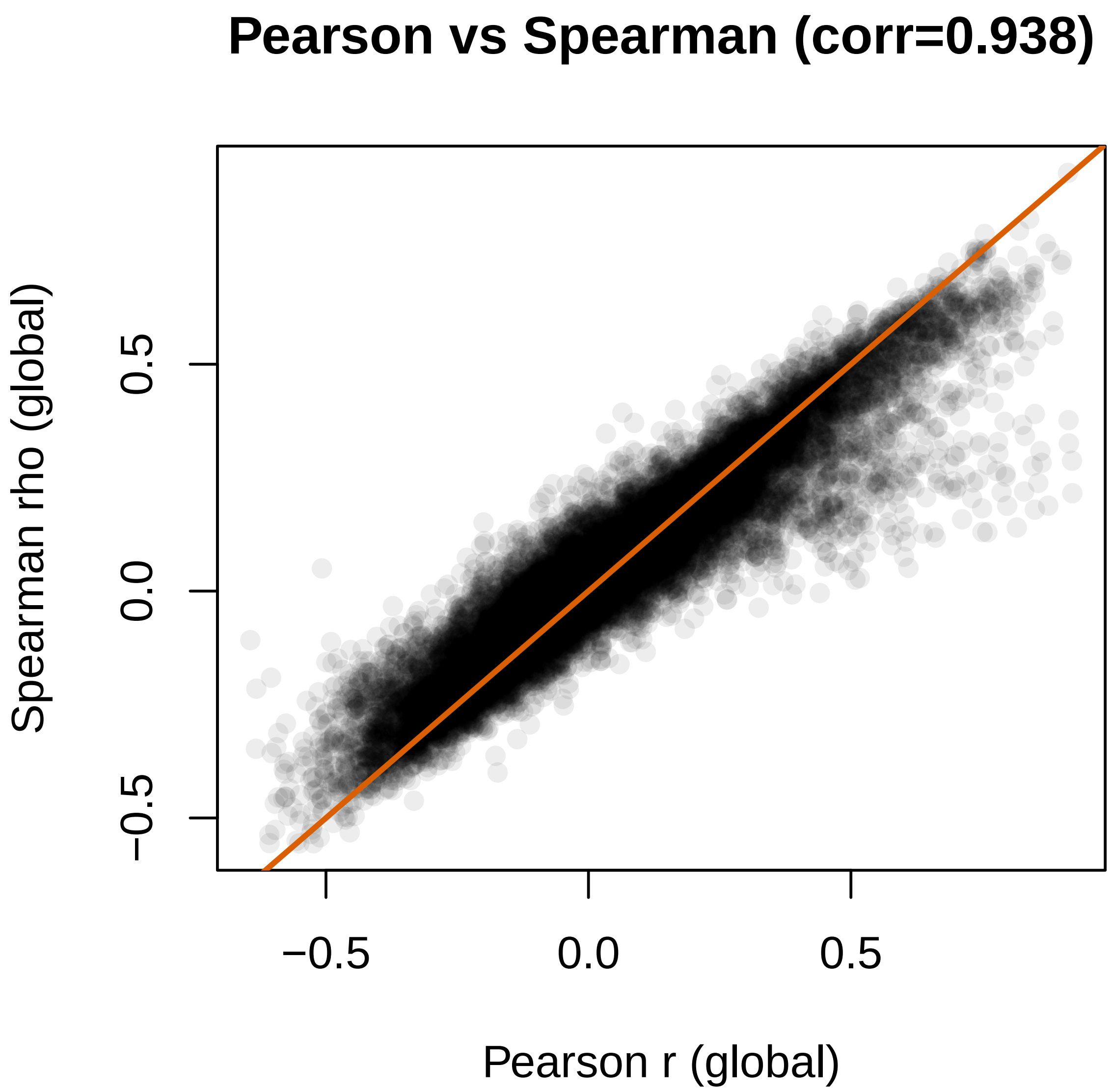


Figure S1: Pearson versus Spearman robustness in TCGA. On a 50,000-pair high-variance subset (1,000 mRNAs $\times$ 50 miRNAs), per-pair Pearson–Spearman agreement averages 0.94 across cohorts. The Simpson-reversal rate at $|r_{\text{global}}| \geq 0.2$ is 12.6% under Spearman versus 13.3% under Pearson, so the qualitative picture does not depend on the correlation measure.

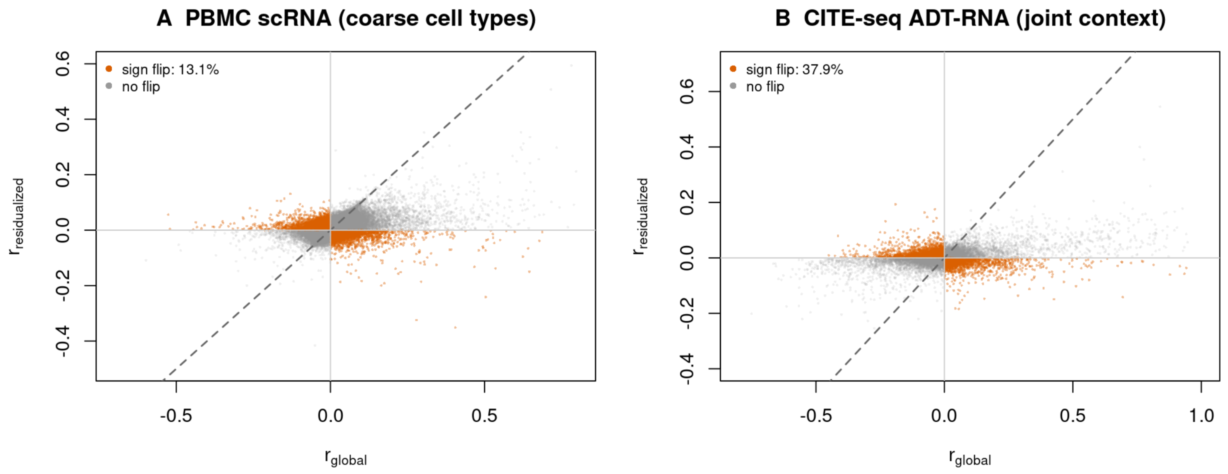


Figure S2: Single-cell global versus residualized correlations across modalities. Each point is a gene–gene pair (PBMC scRNA-seq) or protein–RNA pair (CITE-seq surface protein measured by antibody-derived tag, ADT); $x = r_{\text{global}}$, $y = r_{\text{resid}}$ after cell-type-mean removal. Pairs with $\text{sign}(r_{\text{global}}) \neq \text{sign}(r_{\text{resid}})$ are colored. The diagonal band of flipped pairs at large $|r_{\text{global}}|$ shows that strong pooled associations are not protected from reversal once context is removed.

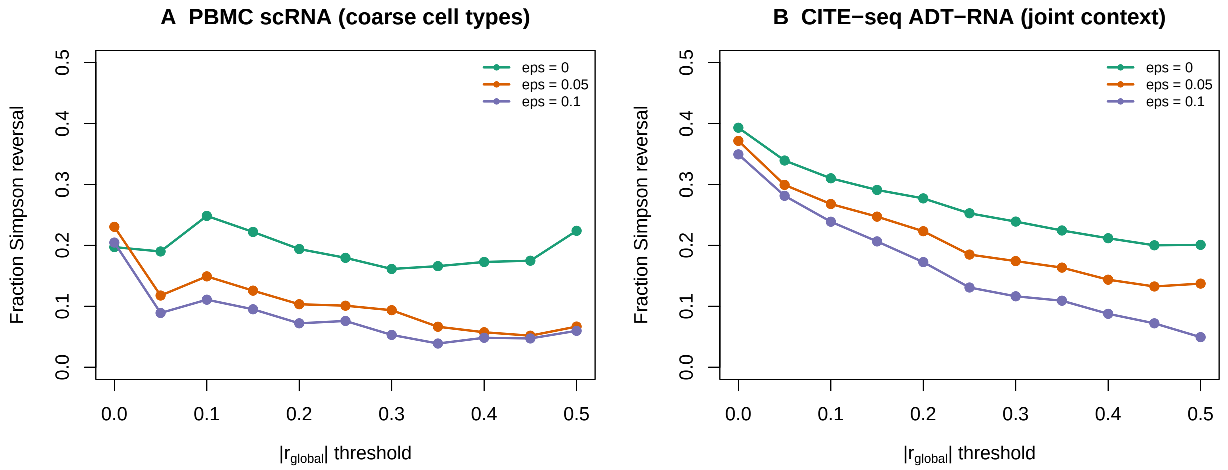


Figure S3: T-cell subtype refinement reduces reversal in PBMC scRNA-seq. Simpson-reversal rate versus effect-size threshold $|r_{\text{global}}|$ for coarse cell types (T, B, NK, monocytes, dendritic, platelets) versus T-cell subtypes (CD4 naive, CD4 memory, CD8, regulatory). At $|r_{\text{global}}| \geq 0.05$ the rate drops from 11.8% to 0.13%; at $|r_{\text{global}}| \geq 0.2$ it drops from 10.3% to 0% (0 of 41 eligible pairs).

#### Supplementary notes

##### Robustness to the correlation measure

The TCGA results are quantified with Pearson $r$, but rank correlation gives the same picture. On a high-variance subset of 50,000 miRNA–mRNA pairs (1,000 mRNAs $\times$ 50 miRNAs), Pearson and Spearman $r$ agree pair-by-pair with mean concordance 0.94 across the 31 cohorts. The Simpson-reversal rate at $|r_{\text{global}}| \geq 0.2$ is 12.6% under Spearman versus 13.3% under Pearson (Fig. S1). Reversal and mixed-sign behavior are properties of the contextual data, not artefacts of distributional assumptions in $r$.

##### Refining cell-type context within PBMCs

Coarse cell types in the PBMC 3k dataset give a 10.3% Simpson-reversal rate at $|r_{\text{global}}| \geq 0.2$. Re-partitioning the T-cell compartment into CD4 naive, CD4 memory, CD8, and regulatory subtypes drops the rate to 0% on the 41 pairs that remain eligible, and from 11.8% to 0.13% at the more permissive $|r_{\text{global}}| \geq 0.05$ threshold (Fig. S3). The same compositional mechanism described in the main text – between-group mean shifts dominating pooled correlations – is what finer partitions absorb: once the dominant mean shift is between T-cell subtypes rather than between T cells and monocytes, the within-context correlations become consistent in sign.

##### Rare stable associations

A small minority of pairs are stably signed across every cohort in which they are measurable. In TCGA, 6,413 of the 692,770 measurable miRTarBase v10 [2] pairs (0.9%) are uniformly negative across cohorts, and the analogous uniformly-signed fraction among strong-effect pairs is well under 1%. These stable pairs include canonical tissue-restricted relationships such as miR-122 [3] within hepatic-lineage cohorts. Their existence confirms that the analysis can detect consistency when it is present, and it sharpens rather than softens the contrast with the heterogeneous majority.

##### Extended methods

Full data-source, preprocessing, correlation, heterogeneity, Simpson-decomposition, mixed-sign, and BRCA-subtype specifications are given in the main-text Methods section. All reproducibility scripts and the "correlation in context" R interface, including the random-effects meta-analytic estimator [4], the Fisher Z transform [5], $I^2$ heterogeneity [1], and Benjamini–Hochberg FDR control [6], are available at `https://github.com/AsiaeeLab/context-corr`.
